# Comment on “Seasonal cycling in the gut microbiome of the Hadza hunter-gatherers of Tanzania”

**DOI:** 10.1101/284513

**Authors:** Stephanie L. Schnorr, Marco Candela, Simone Rampelli, Silvia Turroni, Amanda G. Henry, Alyssa N. Crittenden

## Abstract

In a recent paper, Smits et al. (*Science*, 25 August 2017, p. 802) report on seasonal changes in the gut microbiome of Hadza hunter-gatherers. They argue that seasonal volatility of some bacterial taxa corresponds to seasonal dietary changes. We address the authors’ insufficient reporting of relevant data and problematic areas in their assumptions about Hadza diet that yield inconsistencies in their results and interpretations.

## Main text

To date, robust data has been generated on the gut microbiome of more than a dozen small-scale non-industrial populations practicing varying forms of mixed subsistence, including foraging, farming, horticulture, or pastoralism (1). It is critical for research of this nature, which focuses on biological samples collected from traditionally marginalized groups, many of them indigenous populations, to include an anthropological perspective. This not only allows for the contextualization of dietary and behavioral data, but also acts to inform our understanding of any links to more general conclusions about patterns of human health and disease (1). Microbiome biobanking and large scale data sharing are now a reality (2) and microbes from small-scale non-industrial populations are discussed as a valuable commodity that may be used in the near future for commercial enterprises and health interventions in the post-industrialized west, particularly as therapies for microbiota-driven diseases (3). Therefore, commercial exploitation is a paramount danger, both for vulnerable traditional communities who are the wellspring for microbial biotech inspiration, and for the communities targeted by commercial enterprise with the promise of therapeutic benefits (4). This “unchartered ethical landscape” (5) poses many challenges that require interdisciplinary work between clinical scientists, microbiologists, and anthropologists so that careful attention can be paid to the relevant ethnographic and demographic characteristics of a population under study. As the outcomes of such work often inform strategies of health-cultivation in the cultural west, it is essential that we accurately portray the associative living context of the human populations from which these informative data derive.

Our understanding of how the gut microbiome varies across populations and in response to ecology, subsistence, and health patterns is increasing at a rapid rate. In recent years, several extensive meta-analyses have found that diet composition is one of the drivers of microbiome variation, yet diet in sum may only explain a small fraction of ecosystem diversity found across and within populations (complicated by non-standardized dietary classifications), with often greater mechanistic influence inferable from certain isolated dietary components (6–8). Furthermore, the few studies that have directly explored seasonal variation or short time-series alterations in diet composition have yielded variable results, yet it is important to note that, in each instance detailed diet composition and lifestyle data were collected concomitantly (9–12). Therefore, the effects of temporal change (e.g. seasonality), both short-and long-term duration, on the gut microbiome are not well known and any attempt to investigate the relationship of seasonality on microbiota variation requires detailed supporting data.

In a recent study of seasonal changes in the gut microbiome of Hadza hunter-gatherers, Smits et al. (13) contribute to this burgeoning field by undertaking longitudinal fecal sampling, however, we argue that their characterization of Hadza diet, depiction of seasonal effects, and interpretations of the gut microbiome (of both Hadza and urban industrial populations) are untenable due to insufficient data, inaccurate reporting, and inappropriate analyses. Therefore, we present the following concerns regarding the report by Smits et al.: (1) apparent sample stratification within seasons challenges the conclusions that taxonomic and diversity differences match “seasonally associated functions” rather than specific dietary fluctuations; (2) CAZyme annotations are not provided and CAZyme families are qualitatively summarized using undisclosed criteria (see Figure 2 and Table S9 in Smits et al.), resulting in an incorrect and reductive oversimplification of the array of reported enzymes and the potential microbiome functions (see Table 1); and critically, (3) inaccurate depictions of the Hadza diet throughout the text and a lack of data on diet composition during time of collections (for either Hadza or urban reference sample populations) make evaluation of the results potentially inaccurate and interpretations of seasonal impacts on the microbiome unfeasible.

**Figure 1.**
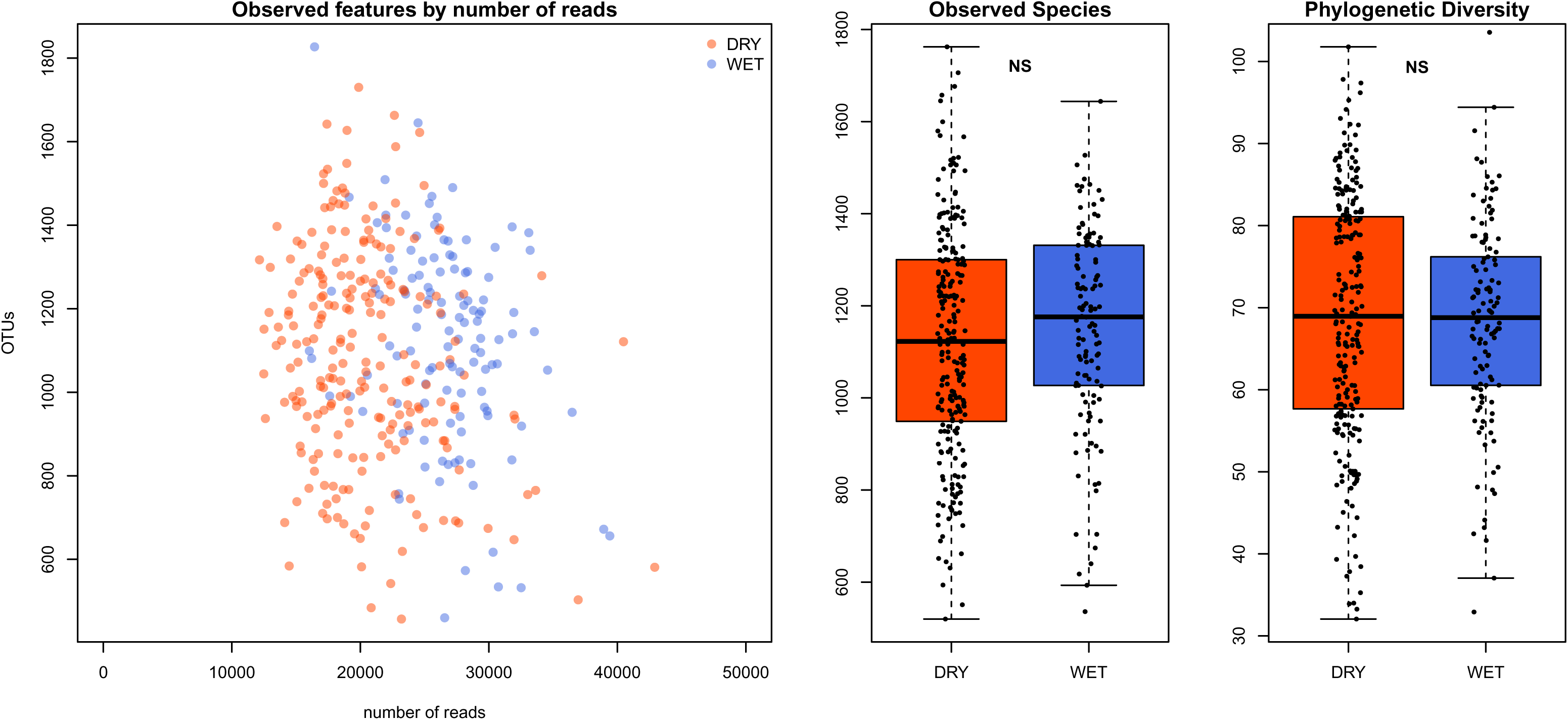
Diversity analysis of 16S rRNA gene data. Reanalysis of the 16S rRNA gene data results in non-significant differences, contrasting reported diversity analysis in Smits et al 2017. Observed features (or OTUs) by read count indicates that wet-season samples in general had greater numbers of reads generated, but with no discernable pattern in OTU diversity. The Observed Species and Phylogenetic Diversity metrics were calculated on OTU tables rarefied to 11,000 reads, and no significant difference is seen between dry and wet season samples based on Wilcoxon test. Samples and plots are colored by season: red = dry; blue = wet.

**Figure 2.**
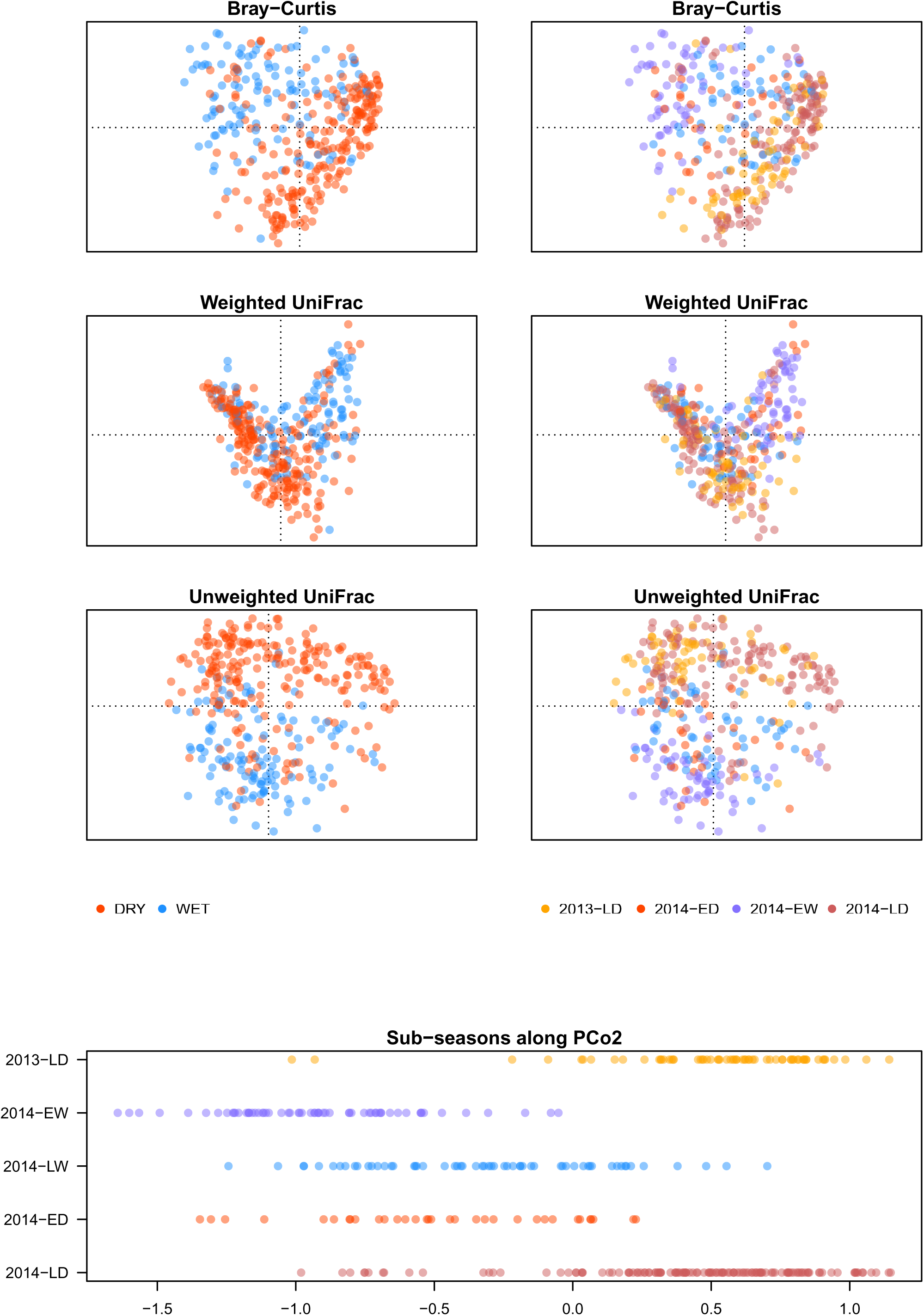
PCoA of 16S rRNA gene data shows stratification by season and sub-season along PCo2 of unweighted UniFrac distance ordination. Ordinations of 16S rRNA gene data show broad seasonal stratification, but sub-seasonal interspersion. A plot of samples along PCo2 of the unweighted UniFrac distance, similar to Figure 1A and 1B of Smits et al., shows sub-season stratification that disrupts clear patterns of seasonal fluctuation. Instead, distribution along PCo2 indicates that the two main clusters apparent in ordination plots above are driven by clustering of the late-dry season data separate from the other sub-seasons. Therefore, seasonal differentiation appears driven only by one sub-seasonal window, which is a time of year when more game meat is available. The late-dry sub-seasons of concurrent years are not significantly varying in their coordinate distribution along PCo2 (p=0.1006), however, significant differences are seen between the early-wet and late-wet sample distribution (p=8.206e-12), between the early-dry and late-dry sample distributions (p=1.065e-13), and between the 2013 dry season samples compared to all 2014 dry season samples (p=5.663e-11). The late-wet season samples and the early-dry season samples do not significantly vary along PCo2 (p=0.147). Dot plots on the left are colored by season (red = dry; blue = wet), and those on the right as well as the bottom panel are colored by sub-season (orange = 2013 late-dry; purple = 2014 early-wet; blue = 2014 late-wet; red = 2014 early-dry; dark red = 2014 late-dry).

**Table 1.**
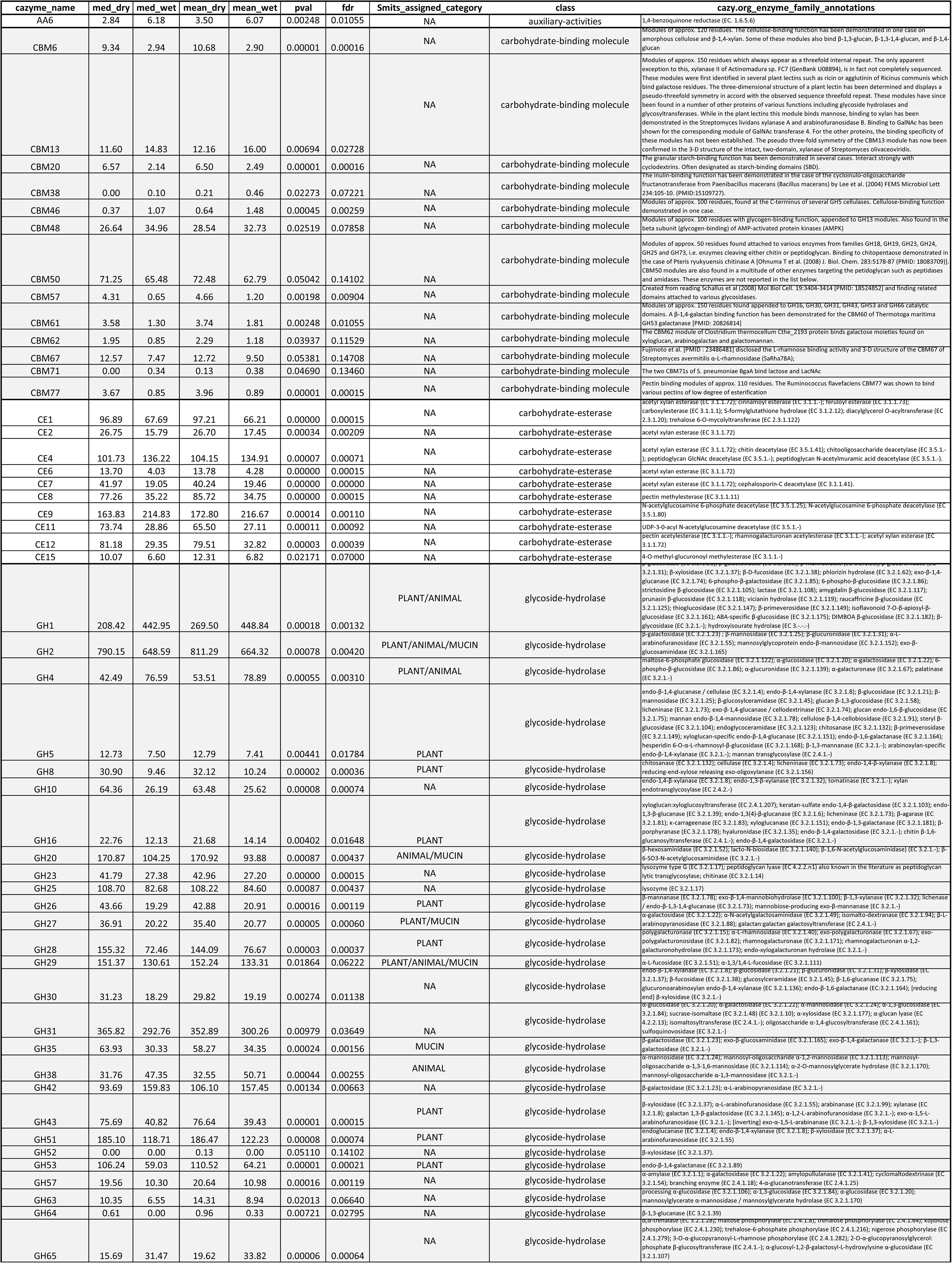
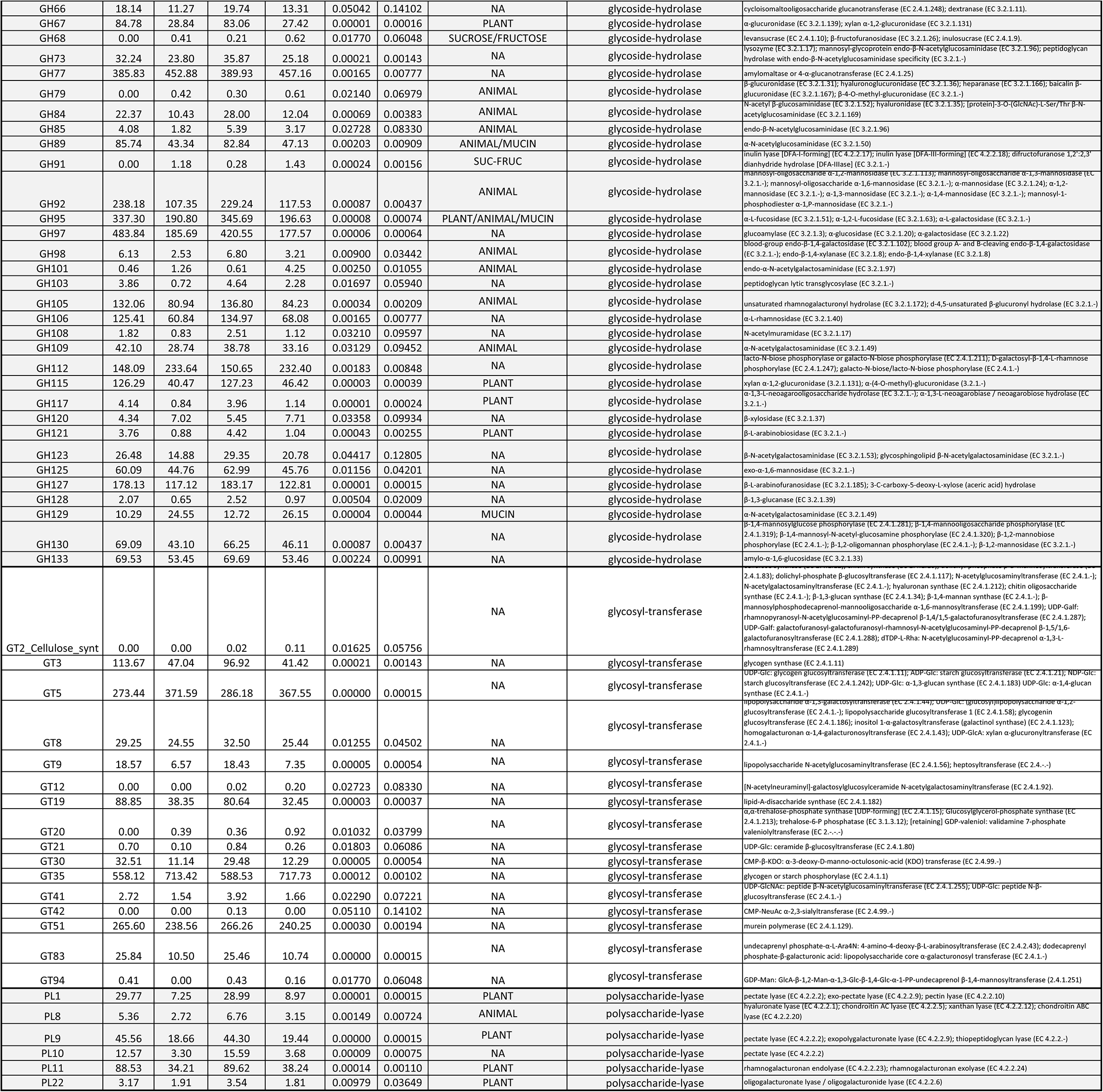
List of the significantly differentially abundance CAZyme families between dry and wet season, determined by Wilcoxon test in R. The median, mean, p-values, and f-values are shown for each CAZyme family. The qualitative category assigned by Smits et al. is provided when available for CAZyme families, along with the actual class type and annotation given from the cazy.org database referencing site.

The Hadza live in a subtropical environment marked by an annual cycle of wet and dry months that determines availability of wild foods. While general trends in food availability are documented in a robust literature going back 60 years, primary production tracks rainfall patterns and often varies annually (14). Smits et al. report higher phylogenetic diversity and observed OTU metrics in dry versus wet season samples and state that, “wet-season microbiotas possess a significantly less diverse CAZyome compared to the dry-season microbiotas,” implying that the more speciose plant-based diet consumed by the Hadza in the wet season (15,16) produces a less diverse taxonomic and CAZyme profile than the meat-rich dry season diet. This finding challenges some prior conclusions about the microbiome-diet relationship that have been made based on observations among post-industrialized populations (11), as well as comparative studies looking at traditional and urban-industrial populations. Thus, the conclusions of Smits et al. contest the hypothesis that more diverse and complex polysaccharide dietary components confer greater gut microbiome diversity (17), which is potentially very interesting and impactful, but requires a sound basis for support. Unfortunately, Smits et al. omit reporting of any observations of dietary intake during sampling dates, do not supply data on rainfall patterns that would corroborate seasonality demarcations during sampling dates, and mischaracterize the prominent seasonal dietary features in their interpretations of CAZyme patterns. It is these concerns that led us to reassess the data reported by Smits and colleagues. The files and R code used to conduct this reanalysis are provided with the supporting documentation.

### (1) Seasonal stratification and diversity

We downloaded the 16S rRNA data from NCBI SRA and conducted data curation and OTU table creation following the same bioinformatic procedures reported in the author’s methods: QIIME 1.9.1 used for quality filtering, OTU clustering with UCLUST at 97% identity against Greengenes 13.8, count table rarefied to 11,000 observations, and diversity analysis accomplished based on the output of the PyNAST aligner and FastTree also implemented in QIIME 1.9.1 (18–20). Our reanalysis does not replicate the finding that taxonomic diversity is significantly greater in dry season samples based on the Observed Species (p=0.1248) and Phylogenetic Diversity (p=0.6448) metrics using a Wilcoxon rank sum test (Figure 1). Similar results were obtained when we used USEARCH UPARSE denovo OTU clustering (results not shown). Importantly, while broad segregation between wet and dry season samples is observed based on ordinations of the Bray-Curtis and UniFrac beta-diversity calculations (Figure 2), interspersion between early and late phases of each season (sub-seasons) is readily apparent. Early and late phases within the same season significantly separate from each other along PCo2 of the unweighted UniFrac PCoA (Figure 2; p=8.206e-12 for wet season, and p=1.065e-13 for dry season), yet the 2014 late-wet and early-dry samples intersperse, and are not significantly varying in their coordinate values (p=0.147; Wilcoxon). These sub-season stratification features, while undisclosed, are nevertheless visible upon close inspection of Smits et al. Figure 1A right panel and 1B bottom panel, and signify stochastic microbiome rearrangements not predicted by seasonality alone. A further problem lies with the early-dry season samples, which were all collected in July 2014, the month of the year when berries often disproportionately contribute to the diet (15,16). Yet Smits et al. characterize dietary trade-offs between wet and dry seasons as follows: “… berry foraging and honey consumption are more frequent during the wet season, whereas hunting is most successful during the dry”. The authors cite Frank Marlowe’s 2010 book as the source of this information, though in the text of this book Marlowe presents data from a previously published study, which found that berries can be a major source of calories during the early dry and the end of the late dry periods (see Fig. 3 of Marlowe and Berbesque, 2009). The specific timing of berry availability depends on rainfall patterns, which can vary significantly from year to year. In order to make accurate statements about the timing of seasons in the East African tropical climate and consequent changes to the diet, it is therefore imperative to record rainfall data during the period of study (14,21). Still, the consistent and cumulative assessments of the Hadza diet across the seasons demonstrates that berry foraging is most prevalent in the early dry season and the end of the late dry season and beginning of the early wet season (15). In this regard, the depiction of Hadza diet seasonality given by Smits et al. is inaccurate with great consequence, as it leads to misunderstandings in further interpretations of their results and a much broader problem for interpreting the drivers in patterns of microbiome taxonomic and functional fluctuations both for Hadza and for the western population samples of the HMP dataset. We must underscore that, despite their incorrect characterization of potential seasonal differences in diet, we nevertheless failed to replicate their reported microbiome patterns for diversity differences and fluctuations in the microbiome that correspond to seasonality.

**Figure 3.**
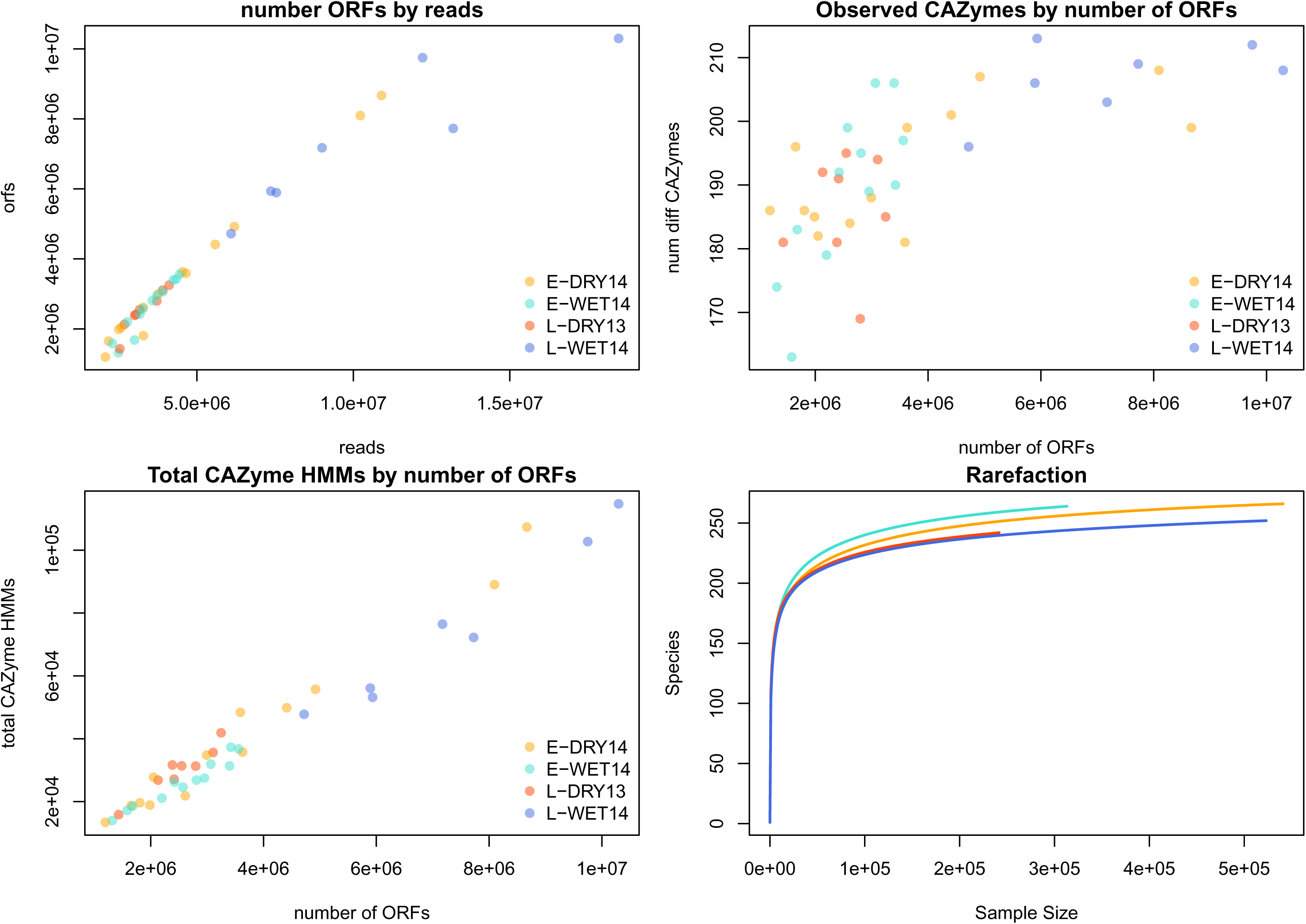
Shotgun data relationships between reads, ORFs, and CAZyme HMM abundances by sub-season. Annotation of ORFs follows a linear pattern based on input read abundance, indicating that ORF prediction follows depth of sequencing, and that CAZyme family HMMs found within each sample scale linearly with the number of ORFs. The number of different observed CAZymes and rarefaction curves indicate a sharp plateau above 200 observations.

### (2) CAZyme annotations and classifications

We reanalyzed the Smits et al. shotgun metagenomic data for CAZymes (22) using hidden Markov model (HMM) profiles on reads, following the same search methods and criteria given in Smits et al. methods: coding sequences, or open reading frames (ORFs), were found from reads with FragGeneScan (23); CAZymes from dbCAN database identified using HMMER (24) with e-value<1e-5, and resultant CAZyme family abundances normalized by the number of ORFs per sample. Descriptive plots of the number of ORFs by number of reads, CAZyme observations and CAZyme hits with sampling estimation curves are presented in Figure 3. Smits et al. report that CAZyme diversity is greater for dry season samples using the Shannon index, which evaluates evenness of abundance across observed “species” (25) and is not necessarily the meaning intuited by “high” or “low” diversity. Shannon diversity is, indeed, greater for dry season samples (p=5.432e-05; Wilcoxon), but the Chao1 measure of richness is equivocal and the “Observed Species” metric is slightly greater for wet season samples (Figure 4). Thus, wet season samples appear to contain greater numbers of different types of CAZyme genes than dry season samples. Conclusions based on differences in overall abundance of CAZyme gene hits depends on whether sub-seasons are lumped or split, whether abundance is normalized by number of ORFs only (Figure 4), or if totals (both rarefied and non-rarefied) are used, indicating potential discrepancies in sampling completeness, bacterial genome size, and gene copy numbers. These details are important because they alter interpretations regarding the size and direction of environmental impacts, and the depth of metagenomic analysis offered by Smits and colleagues does not provide sound conclusions about the adaptiveness of the microbiome functionality, particularly with regards to seasonal variation.

**Figure 4.**
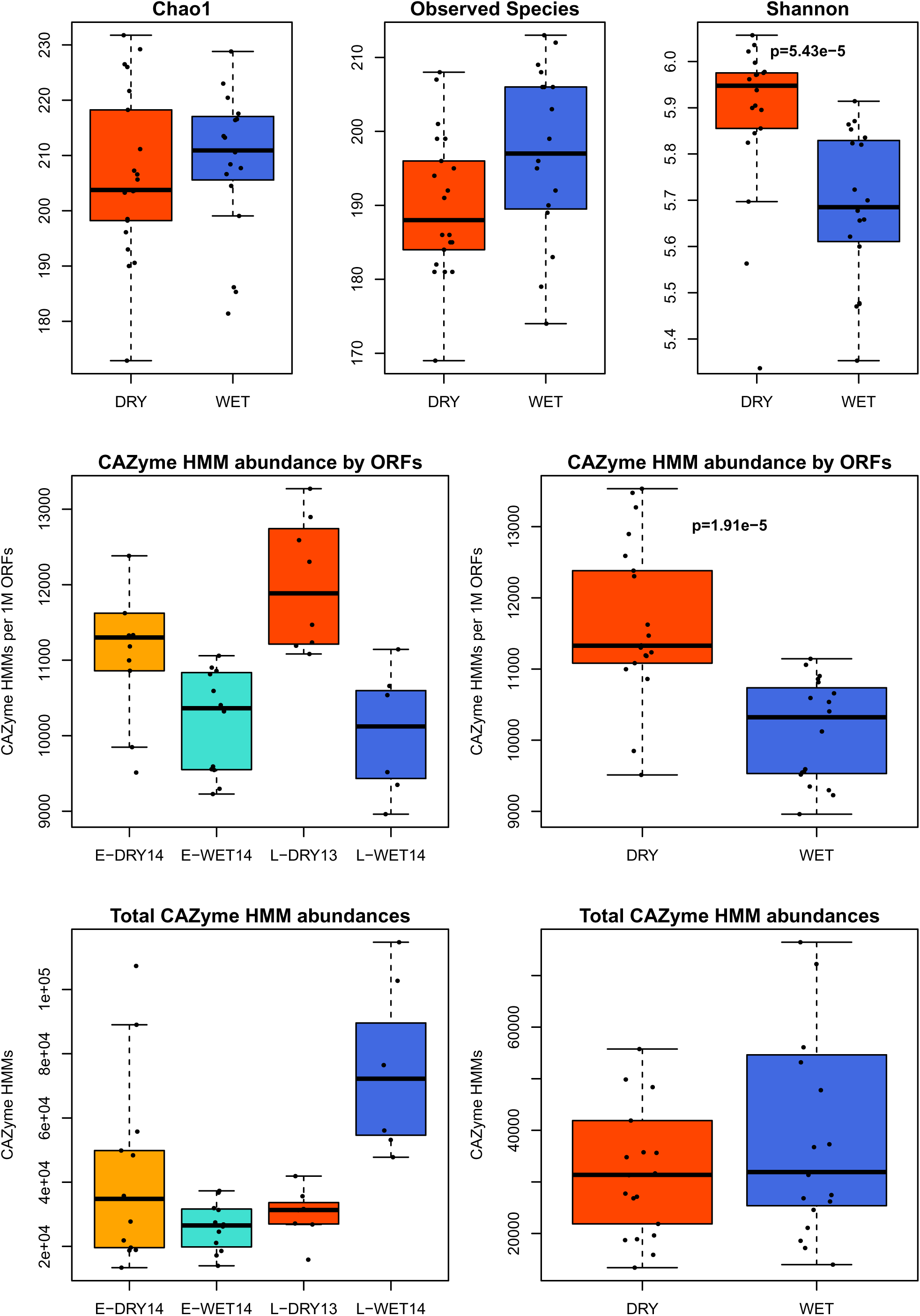
Diversity metrics for CAZyme HMM abundance table and comparisons of total annotations viewed by season and sub-season. Alpha-diversity metrics were calculated on the unrarefied abundance table of CAZyme HMM hits, demonstrating that when other measures of diversity are considered, seasonal differences in diversity are multimodal. Wet season samples appear to have more different types of CAZymes, while dry season samples have a more even pattern of CAZyme abundances. Boxplots of CAZyme abundances normalized per million ORFs clearly show that dry season samples have a larger fraction of the gene-coding regions dedicated to CAZymes, however the late-wet season samples are enriched in total CAZyme HMM counts. This is likely due to read generation being higher for these samples. Gene annotation abundances though must account for genome sizes and gene copy numbers, which vary based on the taxonomic representations in the microbiome, and therefore confound robust interpretations from this simplistic broad-overview analysis strategy. Boxplots colored by season (red = dry; blue = wet), and sub-season (orange = 2014 early-dry; turquoise = 2014 early-wet; red = 2013 late-dry; blue = 2014 late-wet).

An ordination based on Bray-Curtis dissimilarity index and a heatplot of the normalized CAZyme abundances offers an objective view of the stratification of samples based on their profiled functional potential (Figure 5). Again, interspersion between wet and dry season samples is apparent, resulting in three dominant clusters both of individual samples and of CAZymes. The CAZyme families appear alternately abundant in dry and wet seasons and 105 CAZyme families significantly differ in abundance between wet and dry season samples (reduced to 80 after FDR-correction, Wilcoxon; see Table 1). While these clusters could potentially illuminate broad functional associations, it is impractical to ascribe CAZyme families to qualitative categories like “plant”, “animal”, “mucin”, and “sucrose/fructose” because these enzymes may operate in hundreds if not thousands of different pathways and often on secondarily derived molecules rather than the original source substrate (see Table 1 annotations, and see the online CAZy database for a comprehensive list of annotations at http://www.cazy.org/). Additionally, several family categories provided by Smits et al. seem subjectively assigned; many CAZymes were assigned to more than one qualitative category, and so whether a particular CAZyme reflected “plant” or “animal” for a given sample is unclear. As the reported methods do not describe how this process was conducted, the categorization of CAZyme families by Smits et al. is nonreplicable. Another problem is that the CAZyme categorizations given by Smits et al. are misidentified. For example, CAZyme families including chitinases are assigned to the category “plant”, despite chitin being a polysaccharide comprising arthropod exoskeletons and fungi cell walls. Hadza do consume bee larvae from honeycomb, but no particular fungi are known to be a part of their diet. Smits et al. interpret “sucrose/fructose” CAZyme enrichment for wet season samples as associating with berry consumption, which is problematic given data on the timing of berry foraging that we addressed above. It is further perplexing that the authors ignore the overwhelming contribution of honey to the diet during the wet season, which would be a much more parsimonious explanation of “sucrose/fructose” metabolism. Curiously, pectin enzymes are enriched in dry season samples, which is additionally suggestive of either berry consumption or honey consumption having occurred around the time of sample acquisition (August, September, and October) and signifying alterations in typical rainfall patterns (14,16).

**Figure 5.**
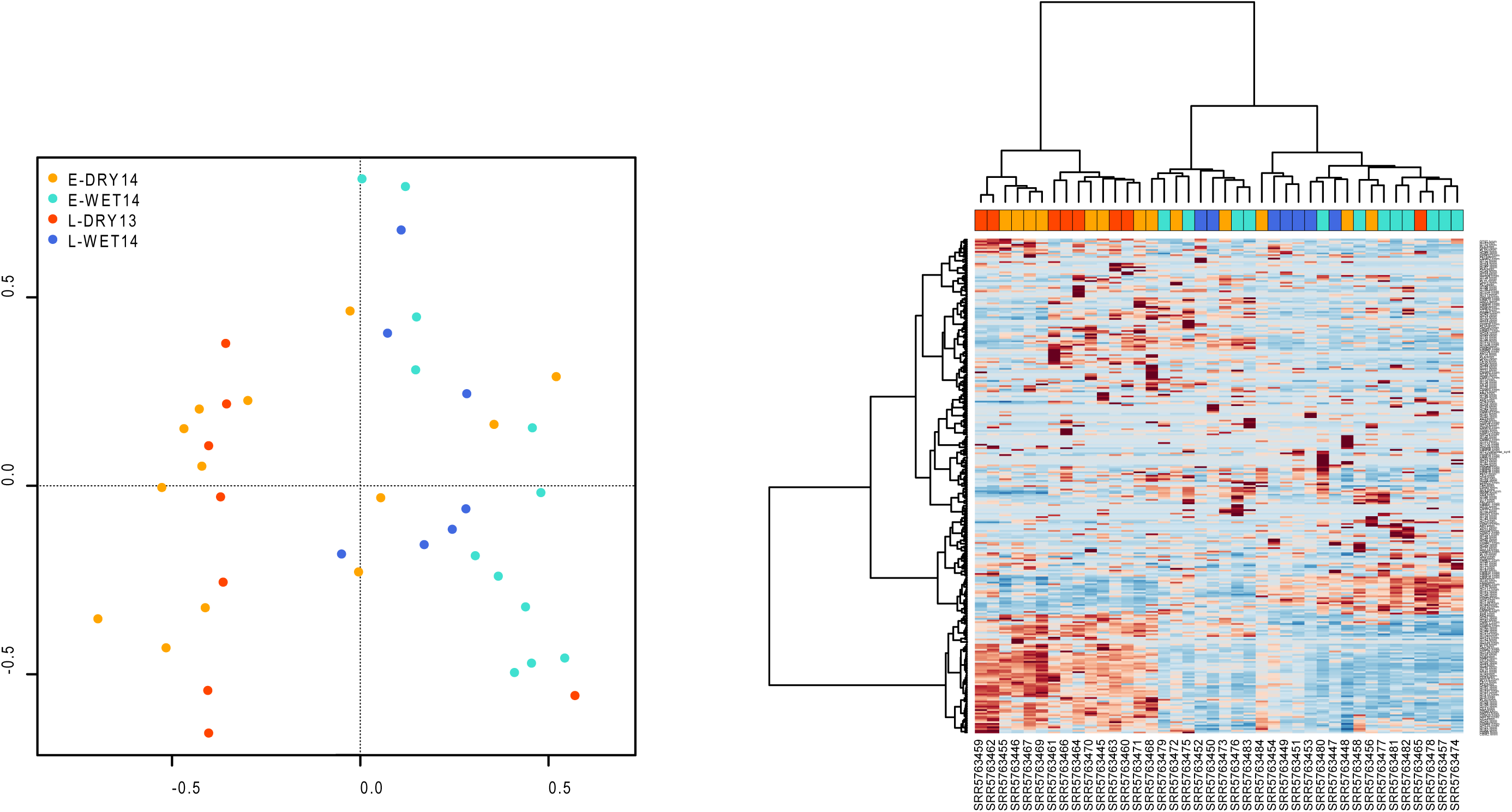
Ordination and heatplot of CAZyme HMM normalized abundance plotted by sub-season shows clustering with interspersion. Ordination using the Bray-Curtis dissimilarity index on the normalized CAZyme HMM abundance table shows variation along PCo1 by wet and dry season samples, with no apparent pattern of distribution based on season or sub-season along PCo2. Heatplot produced from the made4 package in R using Ward’s method for clustering on the normalized CAZyme abundance table with samples in columns, colored by season: orange = early-dry; turquoise = early-wet; red = late-dry; blue = late-wet. CAZyme genes are also clustered in the rows showing three major clusters with sub-clustering of the third. Wet and dry season samples are split along two major clusters, with the wet-season samples sub-clustering and interspersed by some early-dry samples.

The authors go on to state that, “In the dry seasons, the Hadza consume more meat, which corresponds with the enrichment of CAZymes related to animal carbohydrate.” If microbiome fluctuations across the year are a result of dietary seasonal cycling, we would expect the following to be true: greater diversity and abundance of plant foods in the wet season corresponds with enrichment of CAZymes related to plant carbohydrate. However, this logical corollary is not upheld by their results, and acknowledging statements are avoided in the body of the text, but implied in the conclusions. We would expect these reciprocal relationships, at the very least, to be tested and reported. That they are not is a surprising omission, given that the explanation for seasonal cycling posited by Smits et al. rests squarely on seasonal differences in diet, and is a core thesis in their analysis: “Systematic seasonal differences in the Hadza microbiota led us to hypothesize that seasonal dietary changes might lead to related changes in the functional capacity of the microbial community”. Thus, the CAZyme analysis strategy employed by Smits et al. is problematic and distorts conclusions about dietary components that may be contributing to actual observed microbiome features.

Metabolome data reported in Smits et al. Figure S4 are not provided with the manuscript or data files and, therefore, cannot be assessed. However, the KEGG assigned pathways of the metagenomic reads are accessible in a supplementary data file (Table S4) as “relative abundances of genes assigned to KEGG Carbohydrate Metabolism pathways”. We analyzed the KEGG pathway relative abundance data in Table S4 to assess the claim that consistent functional representation is observed across dry season samples, with similar patterns as depicted in the CAZyme analysis. A simple visualization of the KEGG carbohydrate pathway relative abundance data does not yield similar patterns as those depicted in the CAZyme analysis in their Figure 2. In fact, 9 of 15 total KEGG pathways are significantly enriched in wet season samples (Figure 6; FDR < 0.05, Wilcoxon). By plotting the KEGG pathway data provided, we see that dry season samples again intersperse with wet season samples (Figure 7). The relative abundance of carbohydrate pathways are weighted in favor of wet season samples; pathways indicative of plant xylan and hemicellulose metabolism (pentose-phosphate pathway, galactose metabolism), glucose fermentation (glycolysis and pyruvate metabolism), general carbohydrate metabolism of indigestible starch and fiber (propanoate and butanoate metabolism), and metabolism of plant phytates or lecithins (inositol metabolism). This information provides a picture of the seasonal differences in functional potential, whereby wet season samples are actually enriched in carbohydrate metabolism of putatively plant-based carbohydrates (relative to total metabolic function), contrary to what is depicted in Smits et al. Figure 2 and to statements made in the text about analogous patterns of functional enrichments.

**Figure 6.**
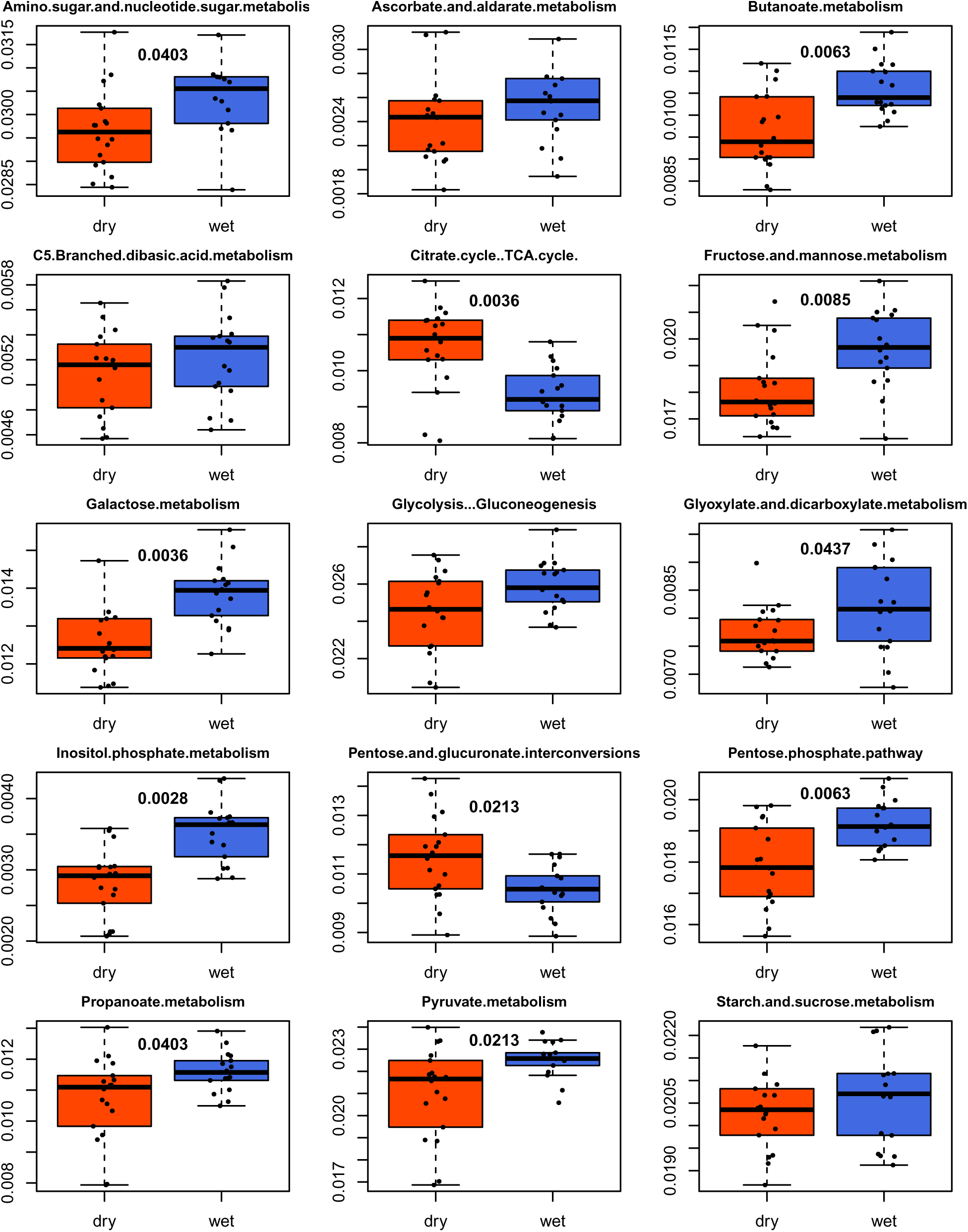
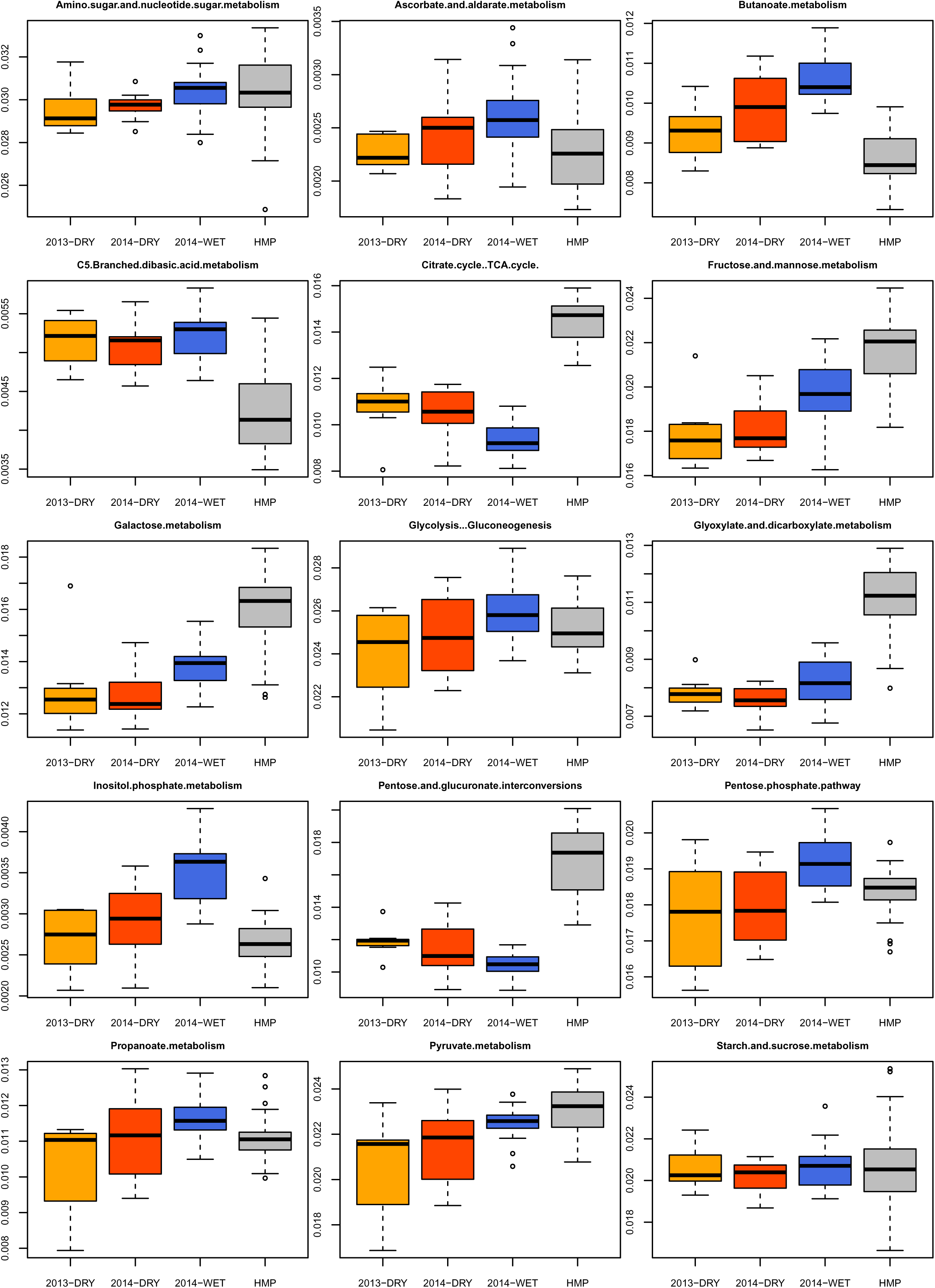
Boxplots for abundance of KEGG pathways in carbohydrate metabolism from Smits et al. Table S4 data trends in favor of wet-season carbohydrate metabolism specialization. KEGG pathway relative abundance supplementary table is visualized with boxplots for only the Hadza samples, colored by season (red = dry; blue = wet). Wilcoxon tests between seasons for each pathway resulted in the finding that 9 of the 15 total pathways are significantly enriched for wet-season samples, with significant p-values displayed in bold in the middle of the plots. These pathways largely correspond with activities for degradation or metabolism of complex polysaccharides and pentose sugars.

**Figure 7.**
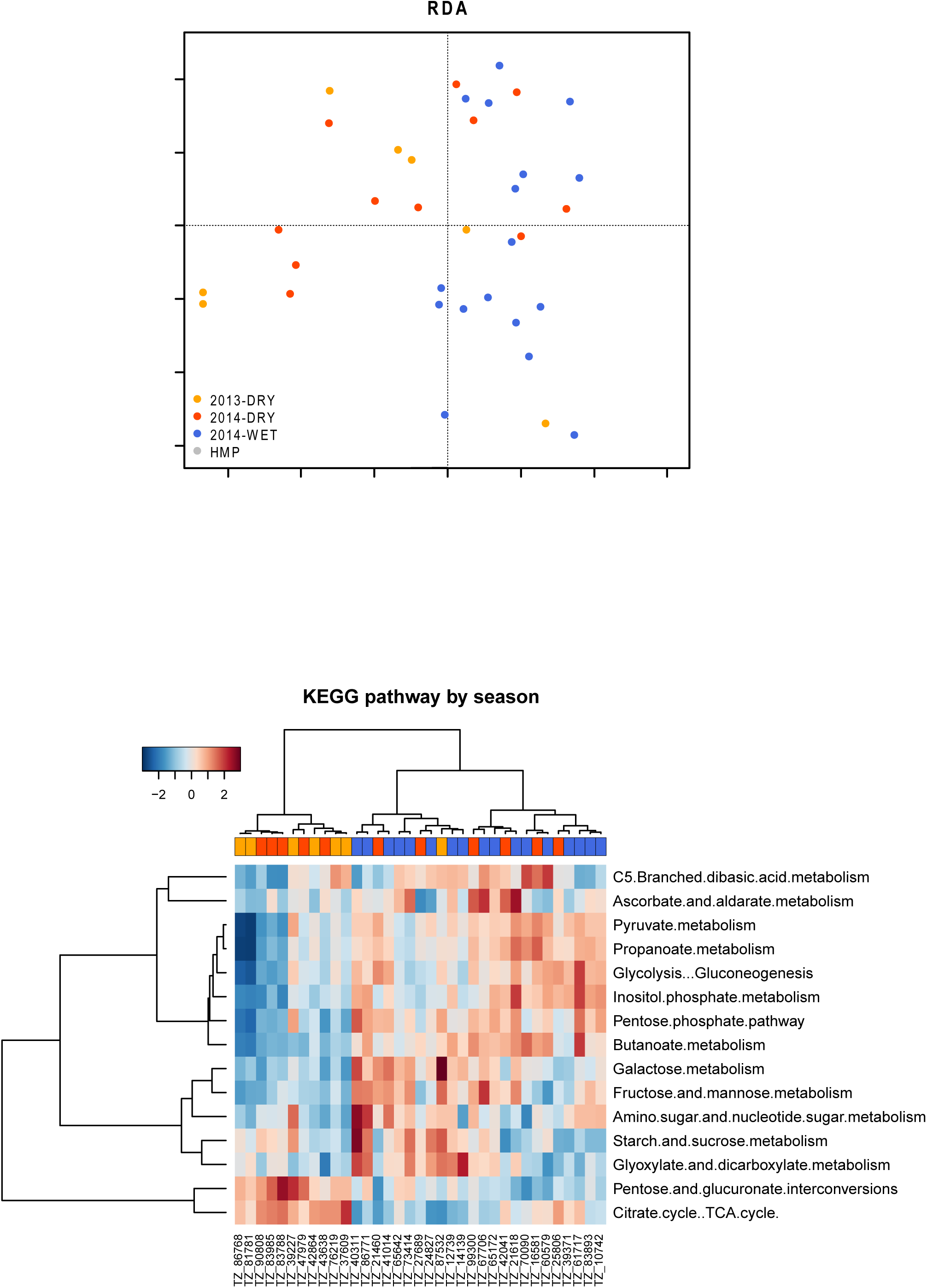
Overview of KEGG pathway abundances for Hadza and HMP samples and ordination plus heatplot of Hadza samples. Broadly, the variation in KEGG pathway relative abundance is greatest between Hadza samples and HMP samples, regardless of season. Hadza samples tend to be inversely enriched in KEGG pathways relative to HMP. Clustering within Hadza samples only shows again separation between wet and dry seasons, but with interspersion. Sub-clustering within the wet season samples indicates trade-off between fiber degradation pathways and starch/simple-sugar metabolism. More KEGG pathways appear enriched in wet season samples than dry season samples.

In general, we find the results and conclusions reported in Smits et al. to be problematic for the reasons outlined above. This extends to the relevant conclusions made at the end of the paper about “seasonally associated functions” and functional association to the “cyclic succession of species”. Neither were seasonal functions unambiguously demonstrated, nor were associations between species fluctuations and gene functions explicitly tested.

### (3) Lack of diet composition data

Perhaps our greatest concern is that dietary information previously indicated as associated with these data were not reported. We would expect, minimally, to see analysis of the “short diet questionnaires for each subject” reportedly collected and analyzed by the team (26). The inclusion of even snapshot dietary composition data at the time of collection could resolve many of the issues outlined here. Since diet is invoked as the main driver for the reported seasonal differences, nutritional information is indispensable. In nearly all instances where population-specific gut microbiome traits are investigated, particularly in studies on seasonality, research teams have provided extensive quantified data on diet (10,12,27–30). Such data are critical when interpreting seasonal impacts on the gut microbiome from populations consuming a seasonally variable diet (10). Furthermore, it is imperative that reported characterizations of diet composition based on findings in the literature are accurately portrayed, and that the dietary conditions under which the data were collected are reported. Only the introduction provides any discussion of seasonal availability of Hadza foods, but this concise summary does neither justice to the complexity of Hadza diet, nor to the robust canon of literature on Hadza diet (see Blurton Jones 2016 for summary of diet related work to date). While we are not suggesting a comprehensive summary of this body of work was in order, we do have reservations that the abridged discussion of seasonality presented by Smits et al. allows for a full interpretation of the data.

The results as presented by Smits et al. posit that taxonomic fluctuations and functional capacities vary by season in a manner contrary to what we might expect given the historical data on Hadza diet and current paradigms about the influence of plant polysaccharides on the gut microbiome, thus their claims should be reinforced by relevant evidence (e.g. dietary and climatic) or include a discussion that addresses discrepancies. Finally, Smits et al. conclude that the loss of seasonal microbiome cycling in western microbiomes may relate to the lack of certain “volatile” microorganisms from the guts of people living in the post-industrialized west. However, the absence of a seasonally sampled correspondent dataset on an urban western population precludes such an inference. Without seasonal environmental or dietary data for the Hadza or for urbanized western controls, it is impossible to interpret what ecological pressures are driving temporal microbiome variation and/or compare microbiome variation in the context of human subsistence. Holding the effects of diet constant, it remains unclear whether or not taxa seen in any population, urban or hunter-gatherer, are, in fact, seasonally volatile.

